# DeepCDpred: Inter-residue Distance and Contact Prediction for Improved Prediction of Protein Structure

**DOI:** 10.1101/425785

**Authors:** Shuangxi Ji, Tŭgçe Oruç, Liam Mead, Muhammad Fayyaz Rehman, Christopher M Thomas, Sam Butterworth, Peter J Winn

## Abstract

Rapid, accurate prediction of protein structure from amino acid sequence would accelerate fields as diverse as drug discovery, synthetic biology and disease diagnosis. Massively improved prediction of protein structures has been driven by improving the prediction of the amino acid residues that contact in their 3D structure. For an average globular protein, around 92% of all residue pairs are non-contacting, therefore accurate prediction of only a small percentage of inter-amino acid distances could increase the number of constraints to guide structure determination. We have trained deep neural networks to predict inter-residue contacts and distances. Distances are predicted with an accuracy better than most contact prediction techniques. Addition of distance constraints improved *de novo* structure predictions for test sets of 158 protein structures, as compared to using the best contact prediction methods alone. Importantly, usage of distance predictions allows the selection of better models from the structure pool without a need for an external model assessment tool. The results also indicate how the accuracy of distance prediction methods might be improved further.

## Introduction

The problem of predicting protein structure from amino acid sequence has been transformed in the last decade from one of aspiration to one of application, although prediction methods are not yet a routine laboratory tool. Recently, well founded predictions of 137 novel folds were published [1]. The authors benchmarked the time for predicting the structure of a 200 amino acid protein as ~ 13 000 CPU core hours, which amounts to around 5 days of processing on 100 cores in a supercomputing cluster, or around 50 to 100 days on a typical desktop machine. This limitation makes structure prediction inaccessible for non-specialists and prevents broader exploitation, e.g. for high-throughput protein structure prediction. Elofsson and co-workers developed a faster high throughput modelling pipeline, but using a less accurate structure prediction protocol, and predicted several hundred novel folds [2]. Here we demonstrate that successful prediction of the distances between residues allows one to predict better structural models. More accurate structures can be generated and better models can be selected from a pool of possible structures than when contact predictions alone are used to constrain the models in the pool. Inter-residue distance predictions thus enhance the ability to generate and select good quality models.

Contacts in a protein structure often involve amino acids that vary across homologs in a correlated way, which is attributed to evolution selecting contacting amino acids to maintain the structural stability of the protein [3] (Fig 1A). However, strong correlation can arise due to two residues contacting a common third amino acid, referred to as a transitive effect, and these residues can thus be falsely predicted as being in contact. During the last decade, efficient global statistical techniques for removing the transitive effect have been developed, thus allowing one to identify clearly the directly coupled positions in multiple sequence alignments (MSAs) [3–7]. For an alignment of length L, such statistical techniques can now predict L/10 contact pairs with accuracies as high as 70 to 80% [8]. A direct result of this has been a rapid improvement in *de novo* structure prediction, e.g. [1, 2], thus fulfilling the original hopes of Valencia and co-workers [9].

**Fig 1.**
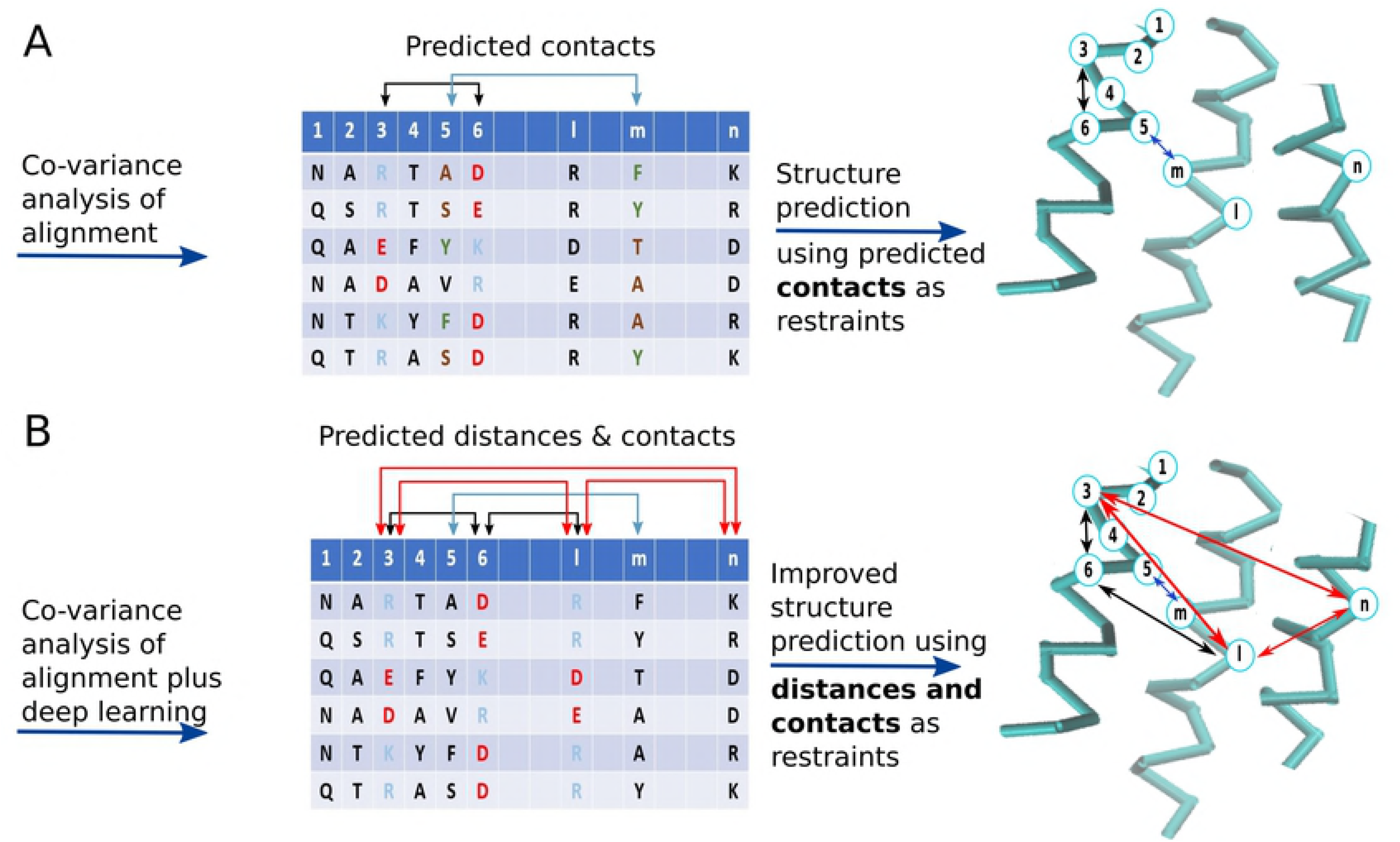
Co-variance between columns of a multiple sequence alignment can be used to predict inter-residue contacts and distances and thence protein structure.

Training neural networks provides further improvement to the accuracy of contact predictions. MetaPSICOV [8] uses a two-layer neural network for contact prediction (i.e. input layer, hidden layer, output layer), with an input vector of 672 features. The feature vector includes local properties of the amino acids under consideration, such as predicted secondary structure and solvent accessibility, properties of the whole sequence, such as the average of the predicted solvent exposure, and coevolutionary scores for directly coupled amino acid positions inferred from the aforementioned global statistical techniques. MetaPSICOV pushes the accuracy of contact prediction to over 90% for the predicted top L/10 contacts, where L in the number of amino acids in the target protein sequence [8]. Very recent applications of deep learning, i.e. multi-layered neural networks, are reported as surpassing the prediction accuracy of MetaPSICOV by 16 to 23% for the top L/5 long range contacts, for the CASP11 protein sets [10–12]. Deep learning increases the number of processing layers so the network can learn to abstract features from the input data, which the final layer can then use for classification. Another recent paper uses a naïve Bayes classifier to calculate the posterior probabilities of eight coevolution analysis, which are then processed by a shallow feed-forward neural network [13].

We hypothesised that pairs of spatially distant amino acids may also co-evolve. Indeed, the literature has some evidence for this [14–16] and Pollastri and co-workers have published distance predictions [17, 18]. However, the RMSD between the actual distance and the predicted distance by this method was over 8 for residues separated by 23 or more residues in sequence. Moreover, *ab initio* models generated with this data had TM-scores of a little over 0.2. TM-score measures the similarity of two protein structures, with 0.2 meaning the predicted structure and the native structure are unrelated. 0.5 indicating that the structures are more likely to be the same fold than not and 0.7 or greater indicating that proteins are almost certainly the same fold [19]. In test systems, adding distance information to contact information could improve structure predictions significantly compared to using contacts alone, even when the distance information was noisy [17, 18]. Thus, accurately predicting precise inter-residue distance from sequence would provide further constraints during structure prediction, with the possibility of increased speed and accuracy (Fig 1B).

We trained four feed-forward neural network models to distinguish residue pairs in spatial distance ranges of 0-8 Å (i.e. contact), 8-13 Å, 13-18 Å and 18-23 Å, which together we call DeepCDpred (deep contact distance prediction). Our method uses a similar feature vector to MetaPSICOV but with eight hidden layers and with only one prediction stage (while MetaPSICOV uses two). The key development is that we predict accurate inter-residue distances, which we show improve structure prediction, and the ability to choose better models from a pool of candidate models.

## Methods

### Neural Network Feature Vector and Set Up

All four distance ranges were trained using the same neural network architecture and inputs, but with training data appropriate to that distance interval. Distances were measured between the C*_β_* atoms (or C*_α_* in the case of glycine) of a pair of residues. There are 733 features as input, described in the supplementary methods, and nine-layer neural network model (i.e. one input layer, one output layer and eight hidden layers where the first layer has 120, second has 50 and the rest have 30 neurons, S1 Fig). For the contact prediction model, networks were trained on distance intervals of (0-7.9]Å, (0-8.0]Å, (0-8.1]Å A and (0-8.2]Å, with the final score for any residue pair being the average output value of all these four. For each of the three distance prediction models, only one neural network was produced.

We implemented here the QUIC algorithm [20] for sparse inverse covariance estimation to calculate amino acid direct couplings for inclusion in the feature vectors. QUIC is similar to PSICOV [5], solving a GLASSO problem, but our tests show that it is much faster than PSICOV, taking on average a quarter of the time to calculate contacts for a 300 amino acid protein, with negligible loss of amino acid contact prediction accuracy (S2 Fig). This allows us to perform calculations on longer amino acid sequences than is achievable with PSICOV.

Residue positions that are very close in protein sequence would be expected to be close in the 3D structure without any need for sophisticated prediction tools. Thus, since they may mask other significant co-evolutionary signals during neural network training, residue pairs separated by 5 or fewer residues were ignored during training and testing, as was also done by others [5–8].

Similarly, it is trivial to predict the distance between two residues on the same secondary structure element, if their sequence separation is known. For the distance predictions, residue pairs on the same predicted alpha helix or beta strand were ignored, to stop their trivial distance prediction; this was done for the training, test and validation sets. In addition, a different minimum sequence separation was set for each distance bin. If the sequence separation of a pair is 5 amino acids, even once residues on the same secondary structure have been ignored, their spatial separation can still be trivially predicted as highly likely to be in distance bin 8-13 Å (S3 Fig). With a sequence separation of 8 amino acids, the distance between them is likely to be in the range of 10-18 Å (S3 Fig, the left blue highlighted bar). Thus, a sequence separation cut-off of 8 or more residues was used for distance bin 8-13 Å and similarly, 13 was the minimum sequence separation for distance bin 13-18 Å. For the 18-23 Å bin a separation of 15 amino acids or more was chosen. For contact predictions, we kept in the data set residue pairs that were predicted to be on the same secondary structure element.

### Training and testing

The main test set consists of 108 from the 150 proteins of the MetaPSICOV test set, so we can be sure that the test proteins are not in the training set of either MetaPSICOV or DeepCDpred. Additionally, these 108 proteins are not listed in the training set of RaptorX [11]. A chain was removed from the MetaPSICOV test set when a sequence with *>*25% identity to it was found in the training set of SPIDER2 [21], since we used SPIDER2 for secondary structure prediction, which is included in our feature vector and for subsequent structural modelling. This gave 108 protein chains ranging from 52 to 266 amino acids with 25% or less sequence identity to each other. Based on annotation in the PDB, 87 chains are monomers in the biological unit, and 21 are from multimeric complexes of some sort, one chain is a membrane protein. The PDB IDs of these 108 protein chains are listed in S2 Table.

Even though the maximum sequence identity is 25% between the training and the test sets, some of the proteins in our test set have common topology classes (and homologous superfamily classes) with the training set proteins, based on CATH classification [22]. In order to test whether our trained model has a bias towards predicting contacts and distances for structures with training set topologies, we generated another test set with 50 proteins that do not have the same topology as any of the training set proteins of DeepCDpred, RaptorX and MetaPSICOV which are listed in S3 Table.

The training set was chosen from the PISCES set [23], downloaded in November 2016. The selected training set protein chains and 50 topologically independent test protein structures were solved with no worse than 2 Å resolution, a maximum R value of 0.25, with no more than 25% pairwise sequence identity to each other or the test set, and with fewer than 400 amino acids. Of these structures, 1701 chains were arbitrarily selected.

The neural network training protocols are described in supplementary methods. The accuracy of the test set predictions was calculated as the true positives divided by the total number of predictions. Since we make no predictions for false positives, i.e. FP=0, the standard formulae for accuracy and precision (PPV) become identical.

### Comparison with other methods

Structure predictions were made using only DeepCDpred contact predictions, using DeepCDpred contact and distance predictions, MetaPSICOV predictions, NeBcon predictions, RaptorX contact predictions and RaptorX contact predictions with DeepCDpred distance predictions. As constraints, the top 3L/2 scoring contacts were used for structure predictions, applying the same Rosetta protocol for all predictions, as described in the next subsection. Residue pairs predicted in the 8-13 Å, 13-18 Å and 18-23 Å distance ranges were selected when they had a neural network score of greater than or equal to 0.6, up to a maximum of 1.5L, L, and L pairs, respectively, for 8-13 Å, 13-18 Å and 18-23 Å bins. For comparison, the RaptorX server was also used for structure predictions.

### Structure prediction protocol

All structures were predicted with AbinitioRelax from Rosetta [24], with constraints applied to enforce predicted secondary structure, contacts and inter-residue distances, as described in Supplementary Methods. Three-residue and nine-residue fragments were created using the program make fragments.pl from the Rosetta suite with the option of excluding homologous structures. We generated 100 candidate structures for each test protein and the one with the lowest total Rosetta energy, including constraint energy, was selected as the prediction, unless otherwise stated. The script for the protocol is given in the supplementary material.

### Determining whether certain residue types are over represented in our predictions

Correctly predicted distances and contacts were examined for biases towards certain residue types. For a score from our neural network of *>*= 0.7, we examined the ratio of the expected distribution (E) in a given distance bin for a given structure, assuming all residue pairs were predicted equally well, to the bias in the distribution that was actually observed (O).

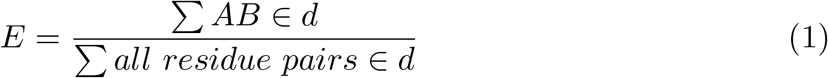

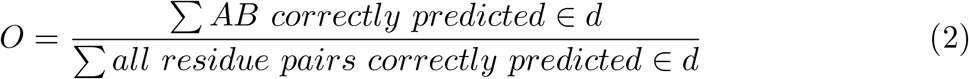

In the above equations, *AB* is a given residue pair type and *d* is the distance bin to which they are assigned based on their spatial separation in the structure under consideration. The mean O/E over all structures was calculated for each pair type, *AB*. We also calculated the fraction of predictions that were true positives for each pair of residue types in each distance bin.

## Results

### Distance predictions lead to improved structure prediction

Our nine-layer neural network was tested on two sets of proteins. The first set tests the network’s ability on the types of proteins that it might encounter in practice, and the second set tests the network’s ability to deal with totally novel folds. The first test set consisted of 108 proteins with 25% or less sequence identity to each other, of which 80 belong to a CATH family homologous to one of the training proteins, 90 have the same CATH topology as a training protein, and the remaining 18 being neither topological nor homologous to our training set. This group represents the sort of sequences that might routinely be submitted to a contact/structure prediction algorithm, 25% sequence identity generally being considered too low for a reliable homology model, even where family homology can be detected [25]. The second test set was 50 proteins topologically different from the training set proteins of our network and of the MetaPSICOV and RaptorX contact prediction neural networks.

The accuracies of the distance predictions for the 108 protein test set are higher than the accuracies of the 50 protein set (Fig 2). For the test set with 108 proteins, distance prediction accuracies are better than the contact prediction accuracies of many other methods. For the test set with 50 proteins, distance prediction accuracies are better than the contact prediction accuracies of MetaPSICOV, but not as high as the RaptorX convolutional neural network (S4 Fig). Comparing with Fig 2 and S4 Fig, the accuracy of the distance predictions falls off much more slowly than the contact predictions, e.g. for the 50 protein set, for the 8-13 Å distance bin, there is a drop of 20 percentage points in accuracy between L/10 predictions and 1.5L predictions, whereas the equivalent drop for contact predictions for RaptorX, DeepCDpred and MetaPSICOV is 40 percentage points.

**Fig 2.**
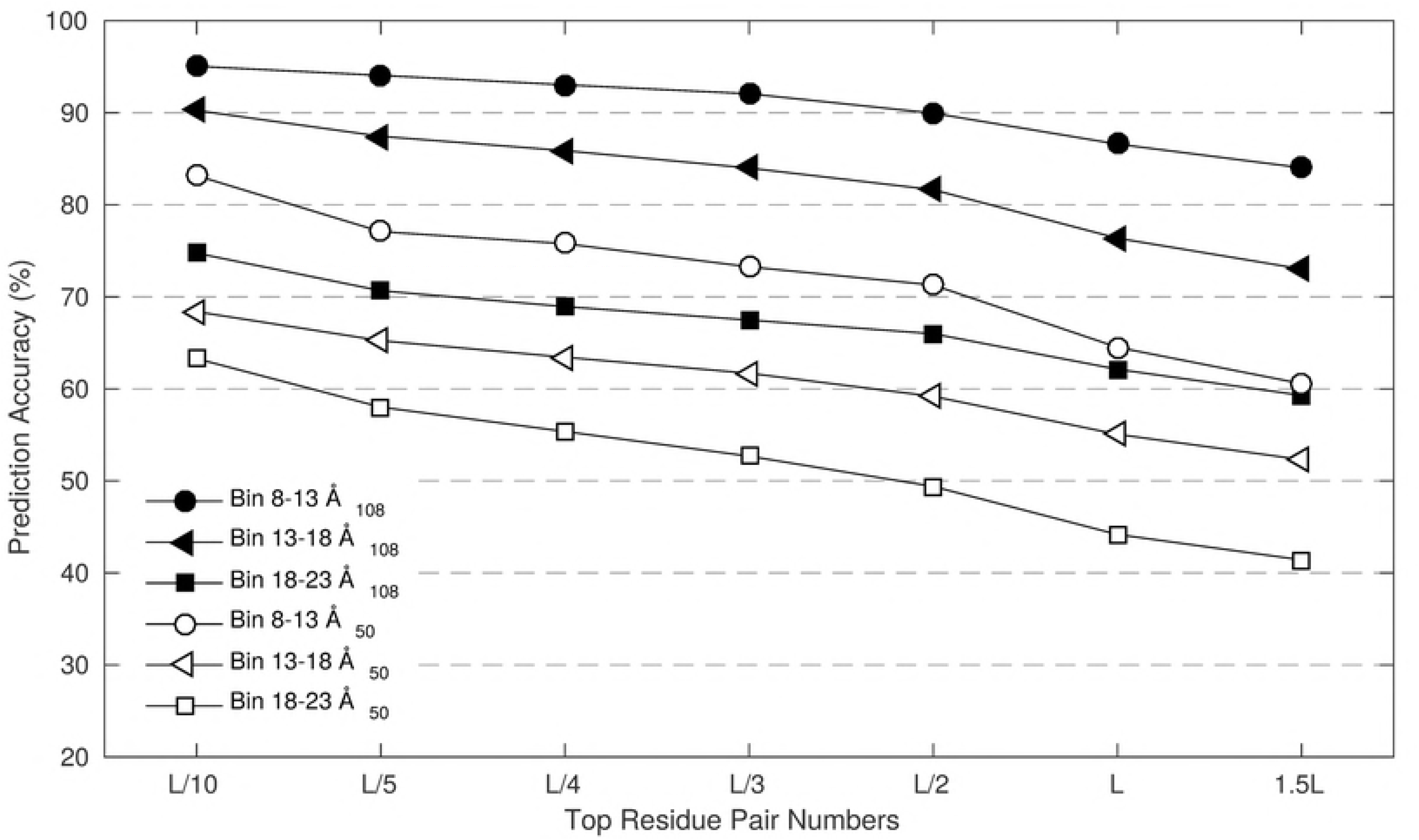
DeepCDpred predicts distances with high average accuracy for both the 108 protein and 50 protein test sets used here.

### Distance predictions lead to improved structure prediction, primarily via better model selection

The model with the lowest Rosetta energy when modelled using distance and contact constraints, from DeepCDpred and RaptorX respectively, is more similar to the experimental structure than when using contact constraints alone. This is true for both the 108 protein test set (Fig 3A) and the set of 50 proteins topologously distinct from the training sets of RaptorX, MetaPSICOV and DeepCDpred (Fig 3C).

**Fig 3.**
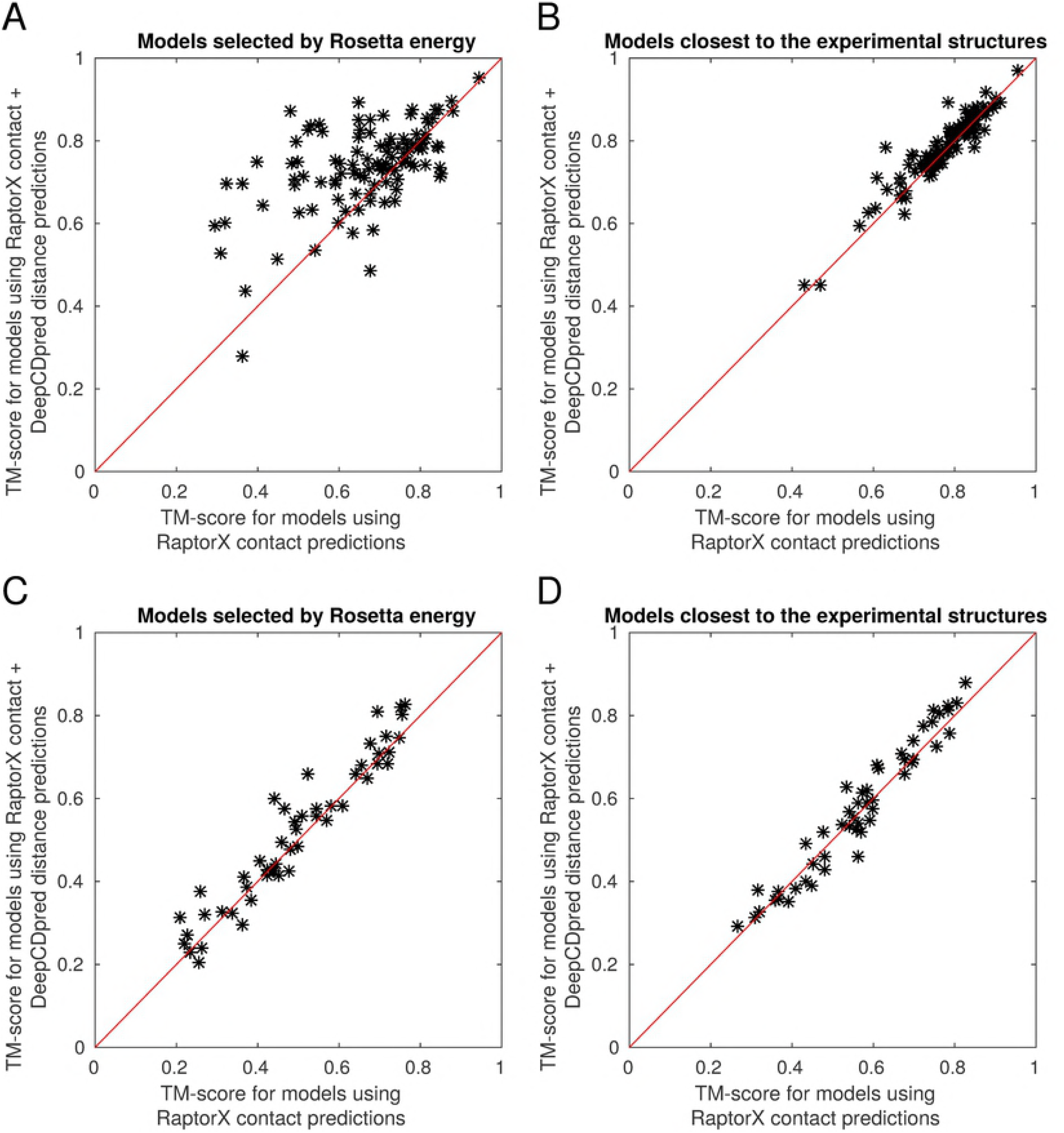
Distance constraints improve structure predictions using Rosetta AbinitioRelax. Addition of distance constraints improves the accuracy of the model with the lowest Rosetta energy (A) and the model closest to the experimental structure (B) for the test set with 108 proteins. For the test set of 50 proteins, distance constraints improve the models with the lowest Rosetta energy (C), but there is small but statistically insignificant improvement in the best model (D). For each test protein 200 structures were generated by Rosetta.

Selecting models by the lowest Rosetta energy, distance constraints in addition to contact constraints improved the mean TM-score compared to experiment by ~ 0.07 in the 108 protein set and ~ 0.03 in the 50 protein set (p-value = 9×10^−9^ and p-value = 0.004 in two paired t-tests, respectively). Inclusion of distance constraints on average improves the best model (i.e. highest TM-score compared to experiment) produced for the 108 protein test set, compared to using constraints only (a p-value of 0.001 in a paired t-test), and has a small but statistically insignificant improvement on the set of 50 proteins (p-value 0.158), with average TM-scores increased by ~ 0.01 and ~ 0.008 respectively (Fig 3B,D).

Thus, applying distance constraints in addition to contact constraints improves the quality of models produced, increasing average TM-scores by ~ 11% and ~ 4% for the 108 and 50 protein sets, respectively, although much of this effect is achieved by the model with the lowest Rosetta energy being close to the best model in the ensemble of models produced by Rosetta, which is not true when contact constraints alone are used (Fig 4). With both test sets ~ 1.4% of the improvement is attributable to improved modelling, with the rest attributable to model selection.

**Fig 4.**
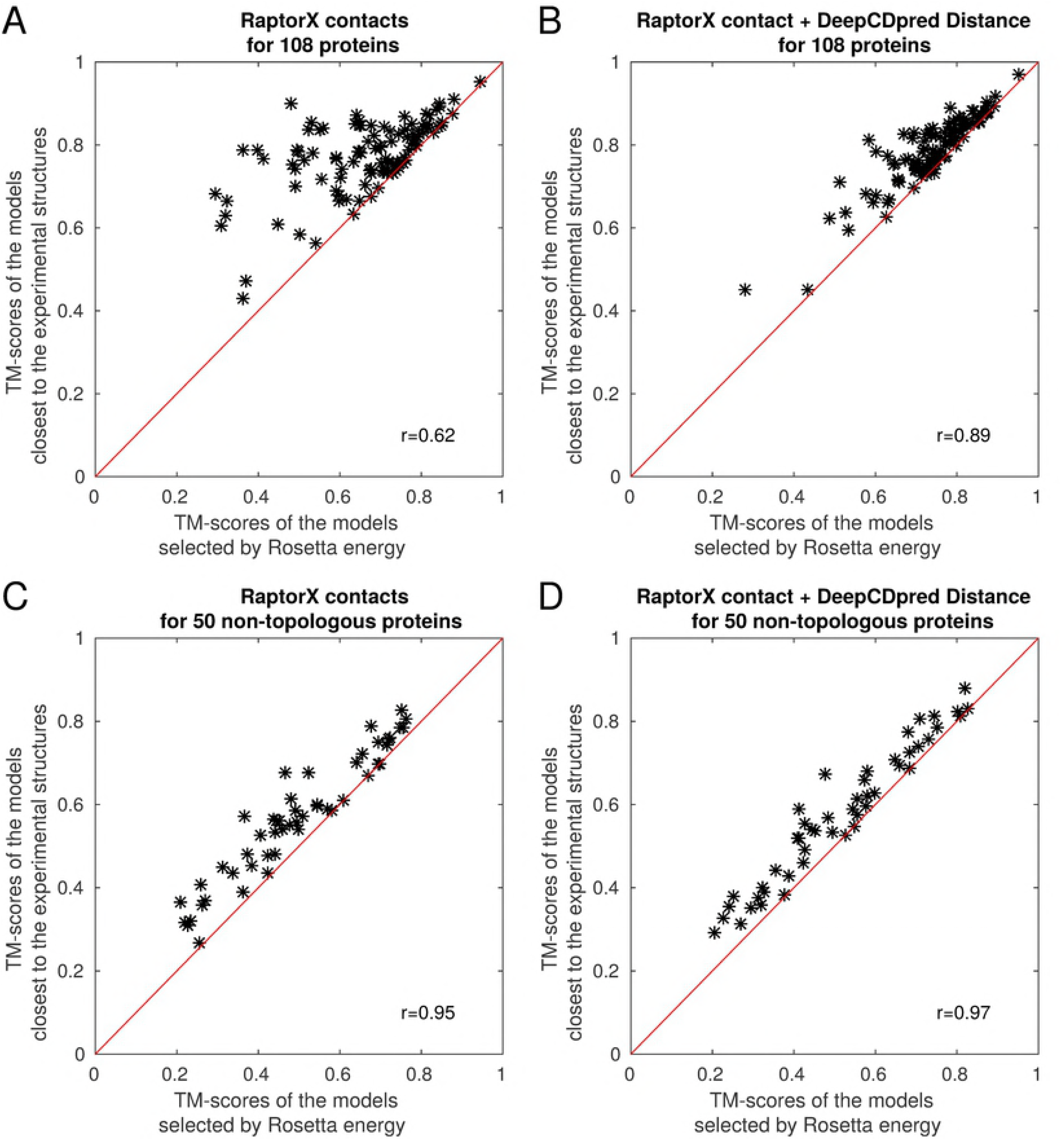
The models with the lowest Rosetta energies, when using distance together with contact constraints, are closer to the best model in the ensemble of models than when using contact constraints alone. For each test protein 200 structures were generated by Rosetta.

### Aliphatic residues are predicted in the contact bins more than expected by chance, but all residue types are equally accurately predicted

To see if any pairs of residue types were disproportionately represented in our predictions, for the 108 protein test set, we analysed all predictions with a neural network score of *>*=0.7. The contact predictions have a higher proportion of hydrophobic interactions than would be expected given the structures under consideration, with aliphatic residues in particular being highly represented (Fig 5). As the inter-residue distance increases, hydrophilic residues become more prevalent in the predictions, but aliphatic residues are over-represented up to a distance of 18 Å, with the 18-23 Å range having hydrophilic interactions disproportionately represented, albeit not as strongly as the 0-8 range is dominated by aliphatic residues. Lysine, and to some degree arginine and glutamate are the most disproportionately represented in the 13-18 and 18-23 Å distance ranges, with valine also continuing to be over-represented. However, the accuracy of a prediction for a given neural network score is largely independent of the residue pair under consideration (S7 Fig).

**Fig 5.**
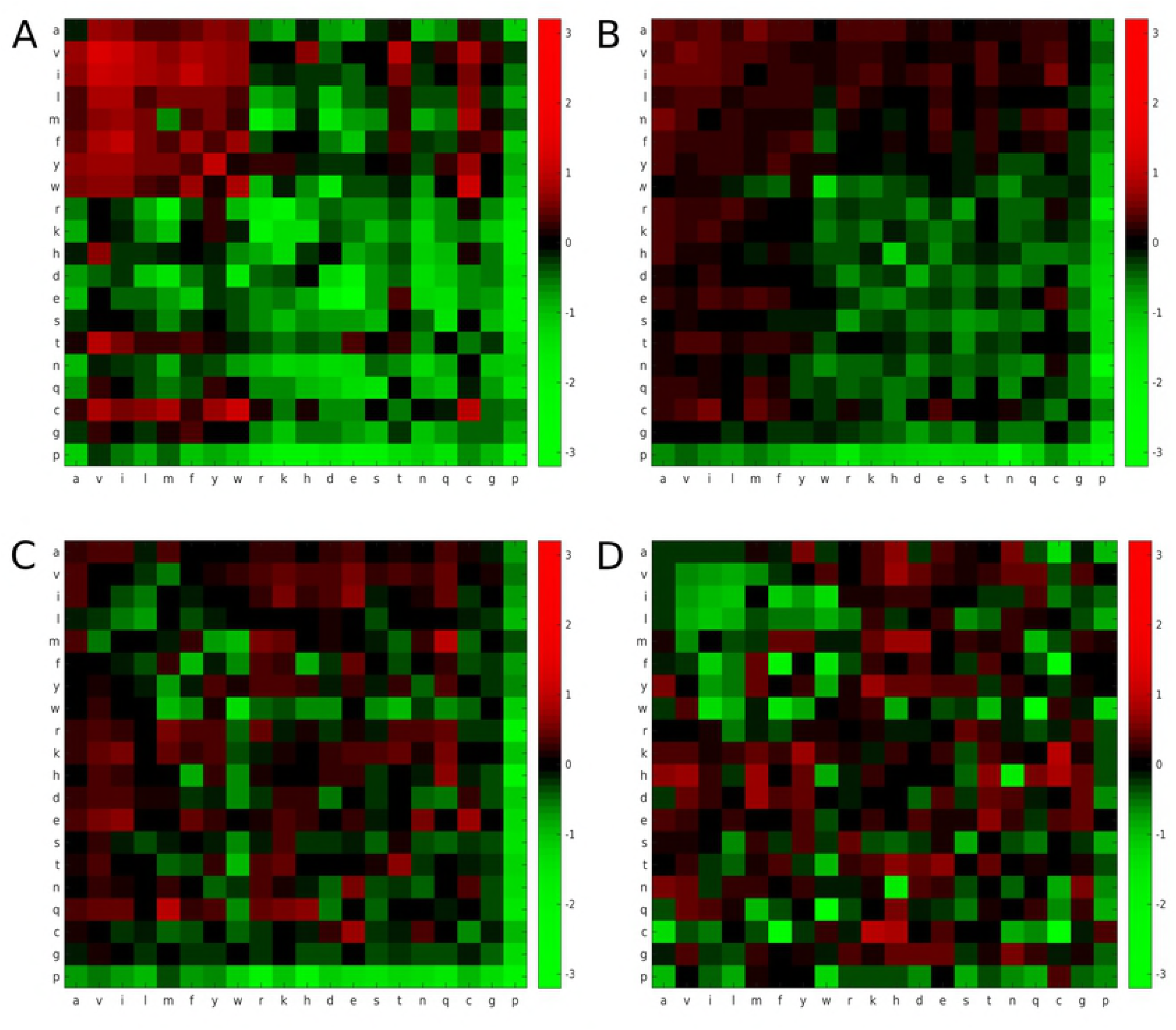
The propensity of correctly predicted contact pairs to be of a given residue type in a given distance bin range. A: 0-8 Å, B: 8-13 Å C: 13-18 Å D: 18-23 Å. Coloured squares represent log_2_(<*O/E*>) as shown on the right hand scale.

## Discussion

There is a large disparity between the accuracy of the predictions of the 108 and 50 protein test sets for distance, contact and structure prediction. In our contact prediction data (S4 Fig), a large part of the difference is due to the set of 50 having much lower Nf values compared to the set of 108 (S9 Fig), where Nf is the number of non-redundant sequences (*M_eff_*) in the MSA divided by the square root of the protein length; *M_eff_* is the number of sequences with *<*80% sequence identity with respect to each other [6]. It has been shown that there is a correlation between the Nf value and the structure prediction accuracy [26]. Sequences were removed from the alignments of the 108 protein set to give them the same Nf as the set of 50, which reduced the accuracy of contact prediction by DeepCDpred, but only accounts for half of the difference between our two test sets (S4 Fig). The accuracy of contacts predicted by the RaptorX server is also affected by reducing the Nf value; the size of this effect for the top 1.5L predictions is approximately a quarter of that seen in the DeepCDpred contact predictions (S4 Fig).

It seems unlikely that DeepCDpred is overfitting the examples, since it was trained with early stopping, and validation and test results give similar prediction accuracies to those of the training data. Nonetheless, the results point to DeepCDpred having poorer generality than RaptorX. Two obvious differences between the prediction methods are that RaptorX uses 6767 training proteins compared to 1701 used for DeepCDpred, and that RaptorX uses a residual neural network, a type of convolutional neural network, whereas DeepCDpred uses a feed-forward network. Increasing the size of the DeepCDpred training set is likely to improve its accuracy for contact and distance prediction. Similarly, investigating the use of alternative neural network architectures, may also lead to improved distance predictions.

It is reasonable to question whether there is a need to pursue improved predictions of inter-residue distance, since one might assume that it is sufficient to be able to predict all contacting residues. It seems unlikely that the goal of predicting all contacting residues will be achievable, since it depends currently on co-variance in residue substitution patterns and there will generally be many positions in a sequence alignment that are totally or highly conserved. The results here demonstrate that using distance prediction can help in model selection and thus improve the prediction of model structure above what can be achieved by the best contact prediction method that was available at the time we undertook this work. Moreover, others report that even in the event of knowledge of all contacts further distance information can improve *de novo* modelling [17, 18]. Knowledge of why the distance between non-contacting residues can be predicted is of interest for trying to improve predictions and also for the potential insight into protein structure and function.

It may be anticipated that the distance prediction could be achieved simply by realising that hydrophilic residues have longer inter-residue distances since they are on the surface and thus in many cases on the far extreme ends of the protein from each other. However, some hydrophilic residues are also in contact with each other on the surface of the protein and thus also form contacts. Constant precision values across all residue pair types (S7 Fig) imply that the fraction of correct predictions is the same irrespective of the residue types. For contacting residue pairs, hydrophobic residues make up a higher proportion of correctly predicted contacts than would be expected based on their frequency of occurrence in the proteins under analysis, i.e. at shorter distance the network has more sensitivity for finding hydrophobic pairs. As the distance increases the network becomes more sensitive for finding hydrophilic residue pairs. It is not clear why there should be this difference in sensitivity, although it may reflect the relative number of examples of the different residue pair types in each distance range, i.e. the network can optimise its error function most easily by becoming more confident at predicting the most abundant residue pair types in a given distance range.

## Conclusion

The data show that inter-residue distances can be predicted reliably using DeepCDpred, the method introduced here. The consequent addition of distance constraints into *de novo* structural modelling leads to better models than when contact predictions alone are used. Including distance constraint terms leads to the models with the lowest Rosetta energy being much closer to the experimental structure than when using only contact constraints together with the Rosetta forcefield. Although others have previously pointed towards the usefulness of distance prediction [17, 18],to our knowledge, this is the first demonstration of the practical benefit of inter-residue distance prediction in the structure prediction problem. We anticipate that improved prediction of inter-residue distance is possible via the most recent developments in deep learning and by understanding the intrinsic bias in amino-acid distribution within protein structures and the effect that has on the accuracy of deep learning methods.

## Supporting information

**S1 Text. Detailed explanation of the implementation** Full details of the feature vector and network architectures for DeepCDpred are explained further here, including the software used for the generation of the feature vector. Details of the structure prediction protocol and a sample from a constraint file are also given.

**S1 Table. Parameters of the contact and distance constraints.**

**S2 Table. PDB ID list of the test set with 108 proteins.**

**S3 Table. PDB ID list of the test set with 50 proteins.**

**S1 Fig. The architecture of the neural network model adopted for amino acid contact and distance predictions in this study.**

**S2 Fig. Contact prediction accuracy and speed comparisons between PSICOV and QUIC.** 221 proteins from the training set were chosen for the comparisons and the accuracies of the top 1.5L amino acid contact predictions of each protein for both PSICOV and QUIC is shown in graph (a). Graph (b) shows the average contact prediction accuracies of the top scoring 1.5L amino acid pairs. (a) and (b) indicate there is little difference between PSICOV and QUIC for amino acid contact prediction. (c), based on the same computer (8-core i7-3770, 32 GB RAM), PSICOV took 16.9 minutes to complete the contact prediction for each protein on average; while QUIC only took 6.9 minutes; especially for large proteins (*>*300 amino acids), QUIC is much faster than PSICOV.

**S3 Fig. The distribution of inter-residue distance with respect to the sequence separation of a pair of residues.** The mean and standard deviation for 435 experimental protein structures from the training set are shown. The three blue highlighted sequence separations (8, 13 and 15) are the minimum sequence separation cut-offs chosen for distance predictions in bin 8-13, 13-18 and 18-23, respectively.

**S4 Fig. Contact prediction accuracies of both test sets (with 108 and 50 proteins).** The average accuracies for the test set with 108 proteins is higher than the test set with 50 proteins. The 108 protein test set had the number of sequences in each MSA reduced to give an average Nf value similar to that of the MSAs for the 50 protein test set. Reducing the Nf value decreased the prediction accuracy of DeepCDpred and RaptorX, however the drop in accuracy of the former was much larger than that of the latter.

**S5 Fig. Addition of distance constraints improves the model quality of both DeepCDpred and RaptorX when the model is selected with Rosetta energy score.** The calculations are for the test set of 108 proteins. The graphs show comparison of the TM-score with respect to experimental structures of lowest energy models predicted using constraints from RaptorX, DeepCDpred contact only, DeepCDpred contact + distance and RaptorX contact + DeepCDpred distance predictions. For each test protein 100 structures were generated by Rosetta.

**S6 Fig. Addition of distance constraints improves the model quality of both DeepCDpred and RaptorX when the model with highest TM-score is selected.** The calculations are for the test set of 108 proteins. The graphs show comparison of the TM-score with respect to experimental structures of the best models predicted using constraints from RaptorX, DeepCDpred contact only, DeepCDpred contact + distance and RaptorX contact + DeepCDpred distance predictions. For each test protein 100 structures were generated by Rosetta.

**S7 Fig. The precision of predicting contacts and distances between different residue types for (a) 0-8 Å, (b) 8-13 Å, (c) 13-18 Å, (d) 18-23 Å.** The scale is given on the right hand side for each plot. Precision is calculated as the number of correctly predicted contacts for that pair of amino acid types divided by the total number of contact predictions for that pair for the predictions with *>*=0.7 network score.

**S8 Fig. TM-scores of the models generated with different tools.** Structure predictions for Rosetta contact and Rosetta contact plus DeepCDpred distances were replicated (replica1 (r1) and replica2 (r2)). For Rosetta server predictions models were selected either by the lowest energy score (CNS score) or the best model among the 5 structures that the server provides. For all other prediction methods, models were selected either with the lowest Rosetta energy or the best TM-score. The calculations were performed for the test set of 108 proteins. The upper and the lower edges of the boxes indicate the 25^*th*^ and 75^*th*^ percentiles, respectively. The medians are shown with the central lines, the means are shown with black ‘+’ signs and the outliers are shown with red ‘+’ signs. Even though the first set of best models which were generated with the restraints of RaptorX contact predictions (RaptorX r1) are significantly better than the best models generated with DeepCDpred contact predictions, replication of the structure predictions with RaptorX contacts (RaptorX r2) resulted in no significantly different average TM-score than the predictions performed with DeepCDpred contacts (paired t-test p-value: 0.507). The results from the RaptorX server were on average worse than all other calculations except the use of MetaPSICOV contact restraints together with Rosetta, presumably because CNS, used by the RaptorX server, is not as good at modelling structures as Rosetta is.

**S9 Fig. Nf value distributions of both test sets (with 108 and 50 proteins).** The upper and the lower edges of the boxes indicate the 25^*th*^ and 75^*th*^ percentiles, respectively. The medians are shown with the central lines, the means are shown with black ‘+’ signs and the outliers are shown with red ‘+’ signs.

**S1 Compressed Folder. Scripts for Rosetta structure prediction and training the neural network.**

## Acknowledgements

SJ, TO, PJW, designed the work; SJ trained and tested 2 and 5 layer networks, implemented the DeepCDpred pipeline and webserver. TO trained 8 hidden layer networks for distance bins, performed predictions for the test set with 50 proteins, performed structure predictions with distance predictions for the test set with 108 proteins, calculated residue pair propensities; LM trained the networks with 8 hidden layers for contact predictions; MFR calculated the contact accuracies of NeBcon for the test set proteins; SJ, TO, PJW wrote the paper, all authors edited the paper and developed project ideas.

